# Modeling Normal Mouse Uterine Contraction and Placental Perfusion with Non-invasive Longitudinal Dynamic Contrast Enhancement MRI

**DOI:** 10.1101/2024.01.31.577398

**Authors:** Devin Raine Everaldo Cortes, Margaret C. Stapleton, Kristina E. Schwab, Dalton West, Noah W. Coulson, Mary Gemmel O’Donnell, Robert W. Powers, Yijen L. Wu

**Affiliations:** Department of Developmental Biology, University of Pittsburgh, Pittsburgh, PA; Department of Bioengineering, Swanson School of Engineering, University of Pittsburgh, Pittsburgh, PA; Rangos Research Center Animal Imaging Core, Children’s Hospital of Pittsburgh, PA; Department of Pediatrics, University of Pittsburgh, Pittsburgh, PA; Department of Biology, Thiel College, Greenville, PA; Magee-Womens Research Institute, Pittsburgh, PA

**Keywords:** **<u>Key Words</u>**: placenta, MRI, perfusion, longitudinal, gadolinium, optic flow, time-frequency analysis

## Abstract

The placenta is a transient organ critical for fetal development. Disruptions of normal placental functions can impact health throughout an individual’s entire life. Although being recognized by the NIH Human Placenta Project as an important organ, the placenta remains understudied, partly because of a lack of non-invasive tools for longitudinally evaluation for key aspects of placental functionalities. Non-invasive imaging that can longitudinally probe murine placental health *in vivo* are critical to understanding placental development throughout pregnancy. We developed advanced imaging processing schemes to establish functional biomarkers for non-invasive longitudinal evaluation of placental development. We developed a dynamic contrast enhancement magnetic resonance imaging (DCE-MRI) pipeline combined with advanced image process methods to model uterine contraction and placental perfusion dynamics. Our novel imaging pipeline uses subcutaneous administration of gadolinium for steepest-slope based perfusion evaluation. This enables non-invasive longitudinal monitoring. Additionally, we advance the placental perfusion chamber paradigm with a novel physiologically-based threshold model for chamber localization and demonstrate spatially varying placental chambers using multiple functional metrics that assess mouse placental development and continuing remodeling throughout gestation. Lastly, using optic flow to quantify placental motions arisen from uterine contractions in conjunction with time-frequency analysis, we demonstrated that the placenta exhibited asymmetric contractile motion.

## Introduction

As identified by the NIH Human Placenta Project, the placenta is an important organ that can impact health across lifespan from conception to the grave. The placenta has a prominent role in fetal growth and development (1, 2) by regulation of nutrient and gas exchange between the developing fetus and the mother. Therefore, placental health directly impacts fetal and pregnancy outcomes (3–5). Furthermore, the placenta not only affects the pregnant person and the developing fetus, but also impacts an array of adult-onset diseases later in life, recognized as the “developmental origins of health and disease” (DOHaD)(6–8). However, its functions are poorly understood, partly because of lacking quantitative and sensitive imaging-based criteria that can continually assess physiological remodeling of placenta throughout gestation.

Placental studies can be difficult to carry out or interpret in human subjects due to ethical concerns of randomization for pregnancy outcomes, wide ranges of genetic backgrounds, and maternal co-morbidity influencing pregnancy outcomes in the singleton pregnancy. Animal models, therefore, are indispensable in modeling human placental conditions for mechanistic understanding and therapeutic development. Murine and human placentas follow similar developmental programs (9–11), but each mouse pregnancy can carry 6-12 fetuses with different genotypes in the same maternal environment, allowing contributions of maternal co-morbidity vs intrinsic fetal factors of the fetoplacental units to be differentiated. With the advancement of gene editing with CRISPR-Cas9 system (12–14), availability of the Mouse Genome Database (MGD)(15, 16), and systematic production of knockout mice with NIH Knockout Mouse Production and Phenotyping Project (KOMP) (17) and International Mouse Phenotyping Consortium (IMPC) (18, 19), an unprecedented number of genetically engineered mouse models are now available for interrogating mechanisms underlying human diseases. Many of these mice have placental abnormalities. A recent screening of more than 100 embryonic lethal mouse mutants found that the placenta is morphologically abnormal in around two-thirds of embryonic lethal mutant mouse lines (1). However, most of the understanding of placental biology came from studying *ex vivo* placental tissues. It lacks a robust and quantitative placental functional evaluation tool for longitudinal tracking of mouse placental development. The objective of our study is to establish MRI-based placental functional biomarkers that can facilitate robust longitudinal evaluation of the placental developmental trajectory in normal wild-type (WT) mice, as a first step towards understanding of placental development. This will facilitate and pave the way for comprehensive phenotyping of mouse models with placental abnormalities.

To support fetal growth and development, the placenta is undergoing constant remodeling and growth. In particular, the labyrinth zone in mouse placenta is the primary location for maternal-fetal nutrient exchange, characterized by an extensive surface area of villi, the finger-like protrusions that contain blood-sinusoids for maternal blood flow. These blood-sinusoids encounter fetal blood vessels and counter-concurrent flow of the two blood supplies facilitates exchange (1–2, 20). The labyrinth zone begins formation around embryonic day 9.5 (E9.5) and is functionally developed by placental maturity at E12.5. However, the labyrinth zone continues to grow all the way through E18.5 (1–3, 20, 21). Additionally, the junctional zone (Jz) serves as the endocrine factory of the placenta. It produces various peptides including growth factors, hormones, and chemokines that facilitate pregnancy progression (20). Most interestingly, it acts on both fetal and maternal physiology and regulates dynamic crosstalk between the decidua and labyrinth zone. The decidua is the maternal tissue that undergoes drastic remodeling after implantation. Specifically, there is large-scale remodeling of maternal vascularization of the decidua. Small blood vessel systems develop into large, dilated ones and there are local changes in vascular elasticity and overall nutrient transport (21). As vasculature remodeling is a critical function of placental development, non-invasive imaging tools to study perfusion, blood-flow, and hemodynamics are critical to understanding placental function as it relates to fetal development and pregnancy outcomes *in vivo*.

We developed a dynamic contrast enhancement MRI (DCE-MRI) protocol that utilizes a subcutaneous administration of gadolinium-based contrast agent (Gd) for the longitudinal visualization of placental development. Our method elicits the ability to conduct longitudinal experimental studies via the ease of subcutaneous administration. This imaging protocol is coupled with advanced image and signal processing biomarkers to understand, monitor, and model placental perfusion, apparent contrast uptake, and fetal motion throughout pregnancy.

We advanced the placental chamber model of the placenta using a novel physiological threshold for the automatic localization. This threshold is derived from stereological analysis performed by Coan et. al. on histological specimens of mouse placentas (22). They utilized grid-based counting methods to evaluate the volume contributions of different placental chambers throughout placental development from E12.5 to E18.5. Our automatic perfusion chamber localization is enabled by applying these volumetric contributions as quantitative thresholds to the perfusion probability density functions of the mouse placenta. We successfully reproduced the paradigm that the labyrinth zone and the decidua can be localized through spatial patterning in peak perfusion and expanded this paradigm to include the steepest slope modeling of the placental perfusion and apparent gadolinium uptake. Differences in the steepest slope represent changes in the instantaneous perfusion rate through the placenta; while the apparent gadolinium uptake measures the cumulative gadolinium in the tissue over the course of the imaging study.

Additionally, we applied computer vision and digital signal processing techniques to study uterine contractions *in vivo*. Phasic uterine contractions are the essential part of parturition for birthing and minimizing postpartum hemorrhage (23–25). Abnormal uterine contractions can lead to labor complications (26, 27) resulting in high risks of maternal mortality and morbidity. It can also cause preterm birth (28, 29), which is the leading cause of infant mortality and morbidity (28, 29). In addition to contractions for labors and uterine atony, the occurrences of Braxton Hicks contractions increase (30, 31) as the pregnancy nears birth. Changes in placental perfusion and placental vascular resistance were observed during the Braxton Hicks contractions (31–34). Although rodent pregnancy with multiple fetuses in the bicornuate uterine horns(35–37) are different from human pregnancy that are mostly with one or two fetuses in the pyriform uteri, rodent models nevertheless are pivotal in investigation of mechanistic understanding of the aberrant uterine contractions to prevent preterm labors or other parturition complications. Phasic contracts of mouse vagina and cervix were observed in *ex vivo* tissues (36). However, it is lacking an *in vivo* tool for evaluating mouse uterine contractions, and how uterine contractions may affect placental perfusion. The two-pronged objective of this study is to establish *in vivo* non-invasive imaging-based biomarkers for quantitative evaluations of mouse uterine contractions in conjunction with placental perfusion.

Briefly, we used optic flow to quantify the placental motion associated with late-term, spontaneous uterine contractions. Further, we characterized the frequency content of the contractile-motion signal via advanced time-frequency analysis: the spectrogram and the periodogram. Thus, we have created the first *in-vivo* research tool for phenotypic investigation of late-term and spontaneous pregnancy contractions. Pre-clinical studies of pharmaceuticals designed to modulate late-term and spontaneous contractions can utilize these methods to longitudinally study drug effects *in-vivo*. Further, basic research can benefit as equally as this work expands the toolbox for longitudinal experimental design. Combining advanced analytical tools with placental MRI optimized for intra-animal temporal studies results in a non-invasive, *in-vivo* pipeline for high-throughput phenotypic studies for key stages of fetal and placental development.

## Results

### Optimization of Subcutaneous Gadolinium (Gd) Contrast Agent Administration for Steepest-Slope Perfusion Modeling

We have optimized the subcutaneous Gd administration and imaging acquisition protocol to enable quantitative placental perfusion mapping with the steepest slope model (38–42). Repetitive intravenous (IV) administration of contrast agent in mice for longitudinal DCE investigation is difficult. Administration via common mouse iv administration routes like femoral, jugular, or retro orbital veins is terminal and cannot be used for longitudinal monitoring. Although tail-vein injection (38, 43–45) can be performed multiple times, quantitative DCE requires administration of Gd contrast agent via an IV catheter inside the magnet during DCE acquisition, which is difficult for multiple administration longitudinally. Therefore, most mouse placental DCE studies with the steepest slope model were performed with single imaging time points. The developmental trajectory across gestation was constructed using different animals, instead of longitudinal monitoring of the same mice. On the contrary, our subcutaneous method can be performed easily with high reproducibility behooving longitudinal studies of disease and fetal development within the same subject. We have optimized the Gd administration timing and concentration to enable the steepest slope model with optimal arterial input function (AIF).

Figure 1A illustrates example DCE time-series images of a mid-slice of a pregnant dam on E14.5 and E17.5 following a single bolus Gd administration via a subcutaneous catheter in the MRI scanner. As Gd contrast agent is taken up by the placenta, the T_1_-weighted signal intensity increases. The multi-planar imaging volume covers the entire abdomen, including the whole litter as well as the maternal kidneys (Figure 1B white arrow) with a clear view of the renal hilum (Fig.1B yellow ellipse) where the arterial input function (AIF) is taken, following established method (58–61). The AIF derived from the kidney hilum, matches that of expected kidney transport dynamics (Fig.1C black). The kidney undergoes rapid “wash-in” uptake of the contrast agent with peak concentration occurring just minutes after injection. Followed by a rapid wash-out phase of the contrast agent in the kidney after nearly 350 seconds after Gd administration. On the other hand, the placental perfusion dynamics (Fig. 1C blue) albeit much slower in their uptake, also illustrate a rapid uptake of contrast agent within the first few minutes after Gd administration, with the steepest slope of the placenta’s temporal concentration gradient occurring roughly at peak AIF following a single bolus Gd administration. After the initial Gd uptake, the placenta continues to take in more contrast agent at a slower rate throughout the rest of the imaging study. The initial placental Gd uptake dynamics, the “wash-in” dynamics, can be quantified by the steepest slope of the MR signal of each voxel.

**Figure 1).**
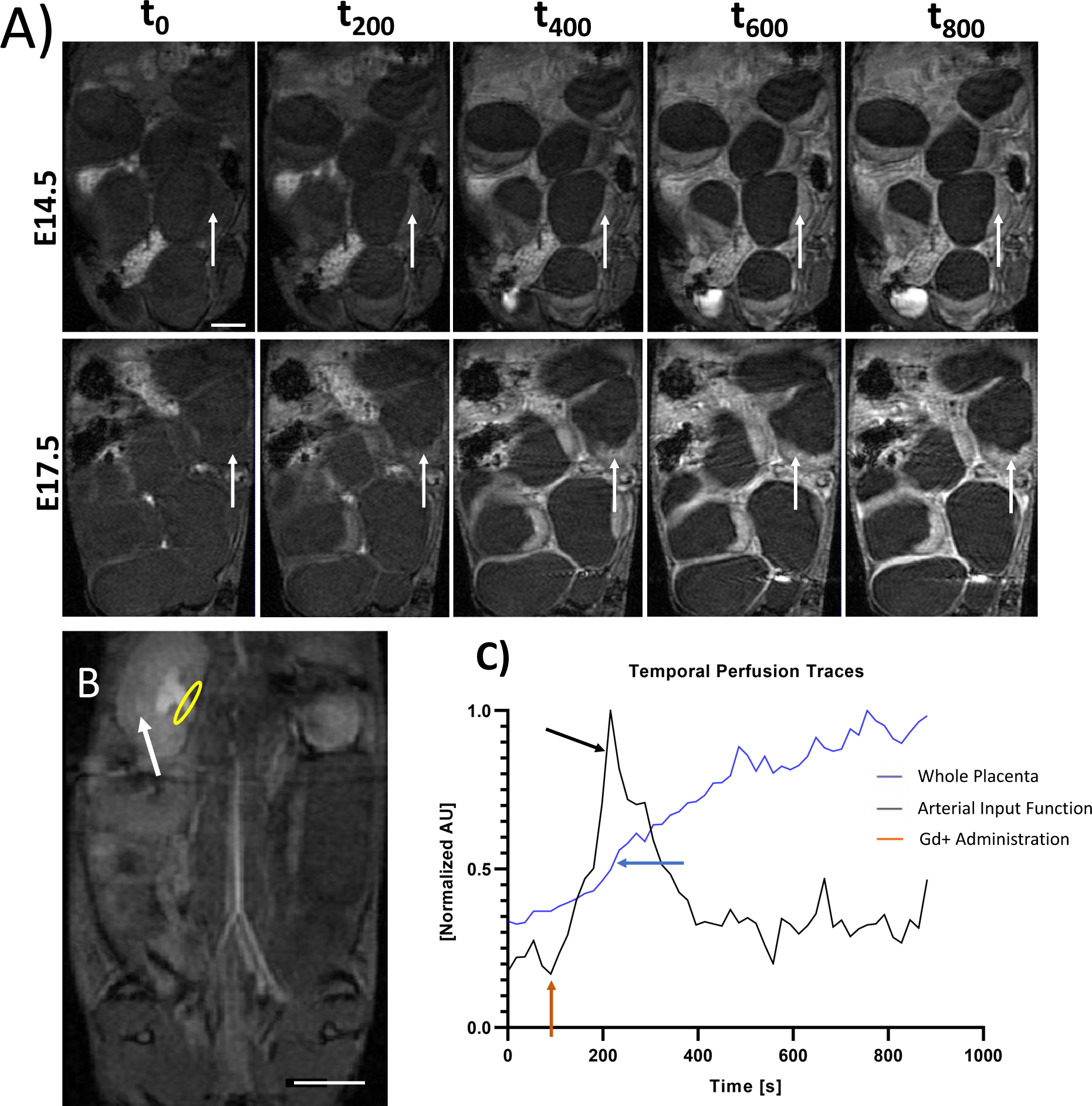
Assessment of SubQ Gd administration for steepest slope perfusion modeling. Example dynamic contrast enhancement time series images for a single slice of a pregnant mouse at A) E14.5 and E17.5 of the same pregnant dam. An example placenta indicated with a white arrow. B) DCE image of the kidney (white arrow) and renal hilum (yellow ellipse) and C) example arterial input functions and whole placenta perfusion trace. The renal hilum is the location from which the arterial input function was determined. Notice in C the sharp increase in kidney perfusion around 200 seconds compared to the steady increase seen in the placenta. Black arrows point the location of the steepest slope of the MR signal time course for the arterial input function (kidney hilum) and placenta. Red arrow indicates the timing of Gd contrast agent administration. The scale bars in A and B are two centimeters.

Voxel-wise steepest slope mapping revealed that the initial Gd uptake dynamics are not spatially homogenous (Fig. 2A, B). There are distinct regions within the placenta that demonstrate faster perfusion dynamics and our physiological threshold successfully localizes these differences into two regions: the high and low perfusion chamber (Fig. 2C, D). The placental development from E14.5 to E17.5 showed differential changes for these two distinct chambers (Fig. 2A-D). The steepest slopes, which reflects the fast initial Gd uptake dynamics, significantly increased (p < 0.0001) in the high perfusion chamber from E14.5 to E17.5 (Fig. 2E)but this was not seen in the low perfusion chamber. When averaging across the entire placenta, the overall steepest slopes were not significantly different between E14.5 and E17.5. The increased steepest slope in the high perfusion chamber from E14.5 to E17.5 is not due to the increase in the AIF obtained from renal hilum. There is no statistically significant difference in AIF between E14.5 and E17.5 (Fig. 2F). This quality control confirms that our observation of the steepest slope increases from E14.5 to E17.5 in the high perfusion chamber are not driven by the global systemic increase of the blood intake dynamic for all visceral organs; but rather by the developmental growth of the placental blood intake dynamic in the high perfusion chamber of the placenta.

**Figure 2).**
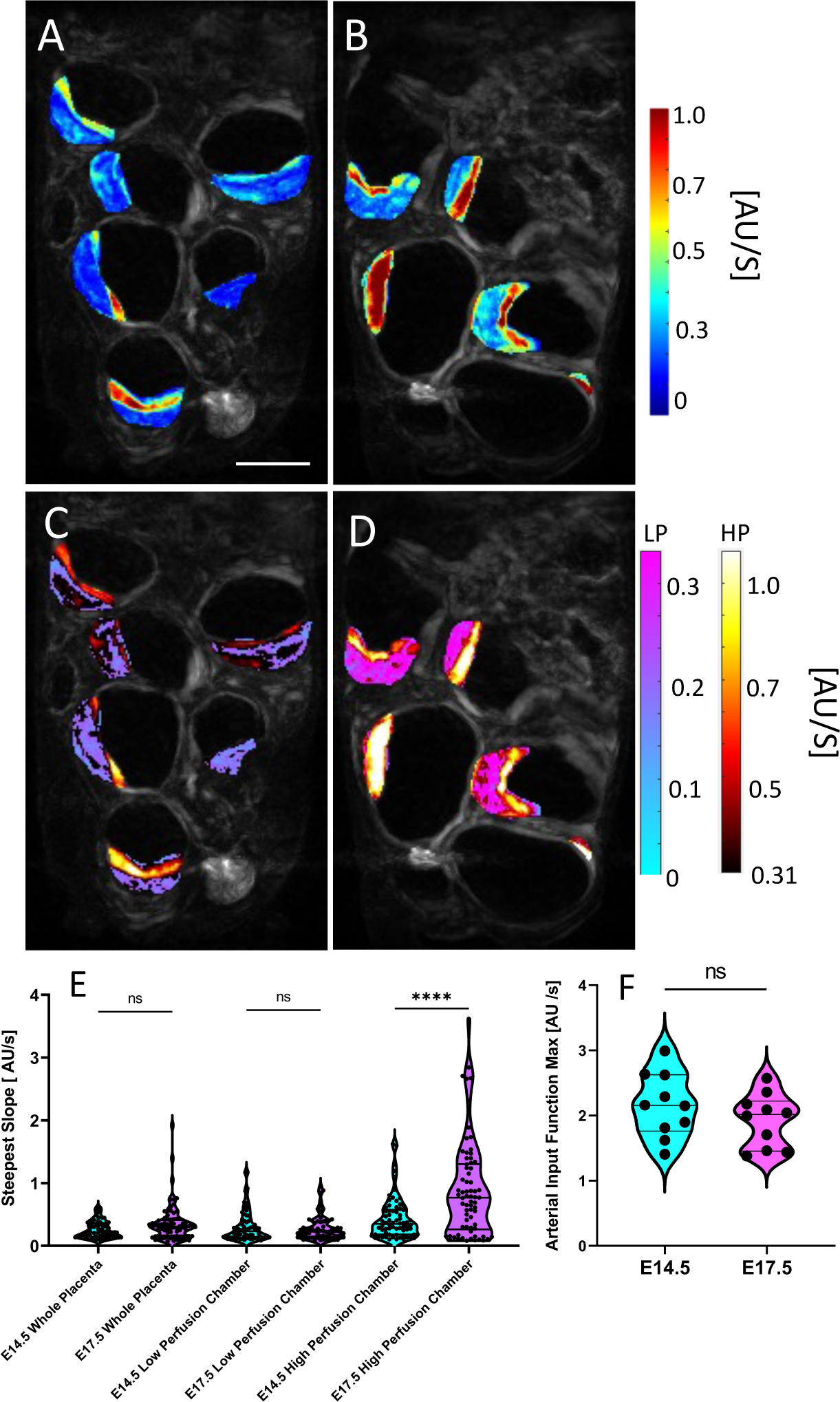
A and B) Steepest slope maps prior to physiological thresholding for E14.5 (A) and E17.5 (B) of the same litter. C and D) Steepest slope maps after physiological thresholding for E14.5 (C) and E17.5 (D). Low perfusion voxels after thresholding use the winter blue-purple colormap while the high perfusion voxels are assigned the black-red colormap. E) Violin plots of the mean steepest slope parameter for the different placental chambers and the whole placenta. Although there is an average increase in the steepest slope parameter for the whole placenta and low perfusion chamber, statistical significance is only achieved for the high perfusion chamber. F) Violin plots of the steepest slope of the arterial input functions for each litter at E14.5 to E17.5. No significant increase in the max of the steepest slope ensures that differences in perfusion are not driven by changes in kidney perfusion dynamics. G) Example arterial input function and placental perfusion trace with the steepest slope of each identified with black arrows. (**** = p < 0.0001). Scale bar is two centimeters.

### Characterization of Placental Development with Quantitative Placental Perfusion MRI

Placental hemodynamic functions have prominent importance for placental health and substrate delivery to the growing fetus. The placental perfusion can be quantitatively mapped (Fig. 3) from the steepest slopes (Fig. 2) in a voxel-wise manner. Figure 3(A-D) shows the voxel-wise perfusion maps of a single slice of the same mouse litter on E14.5 (Fig.3A, C) and then on E17.5 (Fig.3B, D), pre-(Fig.3A, B) and post-(Fig.3C, D) physiological thresholding.

**Figure 3).**
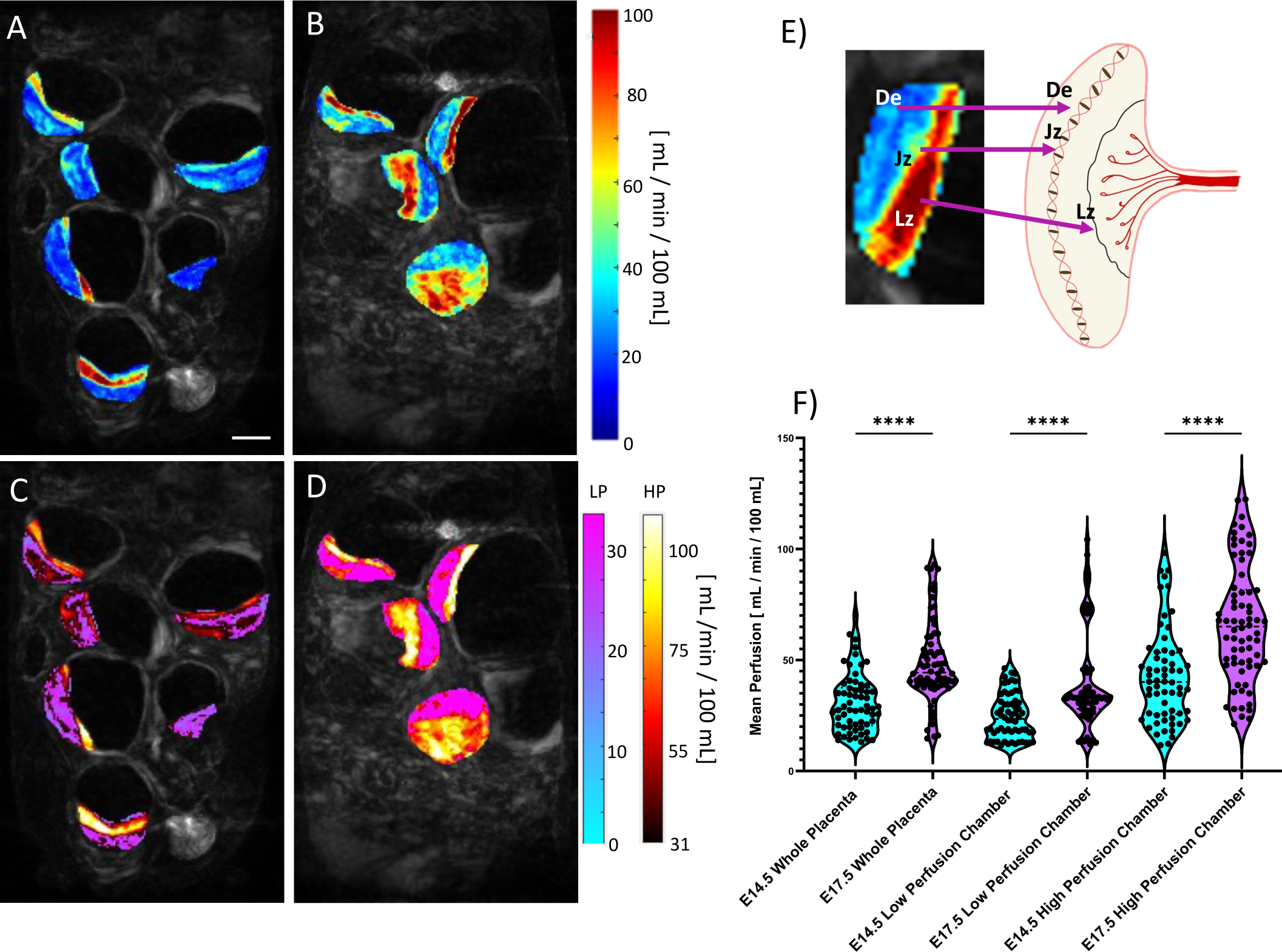
A and B) Perfusion maps prior to physiological thresholding for E14.5 (A) and E17.5 (B) of the same litter. C and D) Perfusion maps after physiological thresholding for E14.5 (C) and E17.5 (D). Physiological thresholding successfully deconvolves the different perfusion zones visually seen in A and B. The low perfusion voxels after thresholding are assigned the blue-purple colormap while the high perfusion voxels are assigned the black-red colormap. E) Perfusion Zones are biologically correlated with different placental chambers. Namely, quantitative perfusion MRI reveals visualization of the Decidua (De), Junctional Zone (Jz), and the Labyrinth Zone (Lz). F) Violin plots of the mean perfusion for the whole placenta and the high and low perfusion chambers at E14.5 and E17.5. There is a statistically significant increase in mean perfusion in the high perfusion chamber from E14.5 to E17.5. (**** = p <0.0001). Scale bar is two centimeters.

On average there is an increase in total placental perfusion. The low perfusion chamber average perfusion increases by ∼15.6 mL/min/100 mL from E14.5 to E17.5, whereas the high perfusion chamber average perfusion increases by ∼24.97 mL/min/100 mL (Fig 3F). The peak perfusion in each chamber from E14.5 to E17.5 was statistically significant (p < 0.0001). The anatomical locations of the high and low perfusion chambers correlate with known placental anatomy (Fig. 3E): the high perfusion chamber correlates with the labyrinth zone whereas the low perfusion chamber correlates with the decidua, separated by a strip of intermediate perfusion zone, correlated with the junction zone (Figure 3E). The high and low perfusion chambers isolated by our physiological thresholding model corresponded well with the location, size, and morphology of a healthy mouse placenta on E17.5 (22).

Longitudinal perfusion analysis can be performed at the litter-level or on the individual placentas. Figure 4 A, B details average litter-wise (Fig. 4A) and placenta-wise (Fig. 4B) longitudinal analysis of the mean perfusion for the whole placenta, low perfusion chamber, and the high perfusion chamber. Placental tracking from E14.5 to E17.5 was done through visual inspection of the DCE-MRI time series, spatial location, orientation and size of the placenta were used to indicate a likely pairing. There is on average an increase in the mean perfusion within the high perfusion chamber for every placenta from E14.5 to E17.5. The slope of the linear best fit (Fig. 4A, B) for litter wise analysis of the low, high perfusion chambers, as well as the whole placenta is 5.23 ± 2.67 mL/min/100 mL, 8.23 ± 3.15 mL/min/100mL, and 5.83 ± 2.23 mL/min/ 100 mL, respectively. The placenta wise linear best-fit (Fig. 4B), has similar slopes to the litter wise fitting with 5.13 ± 0.97 mL/min/100 mL, 7.82 ± 1.2 mL/min/100mL and 5.64 ± 0.89 mL/min/100 mL for the low perfusion, high perfusion chamber, and whole placenta, respectively. The litter-wise linear best-fit and 95% confidence intervals (Fig 4A, B) contains much more overlap than placenta-wise analysis, indicating individual litters show variation in perfusion magnitude.

**Figure 4).**
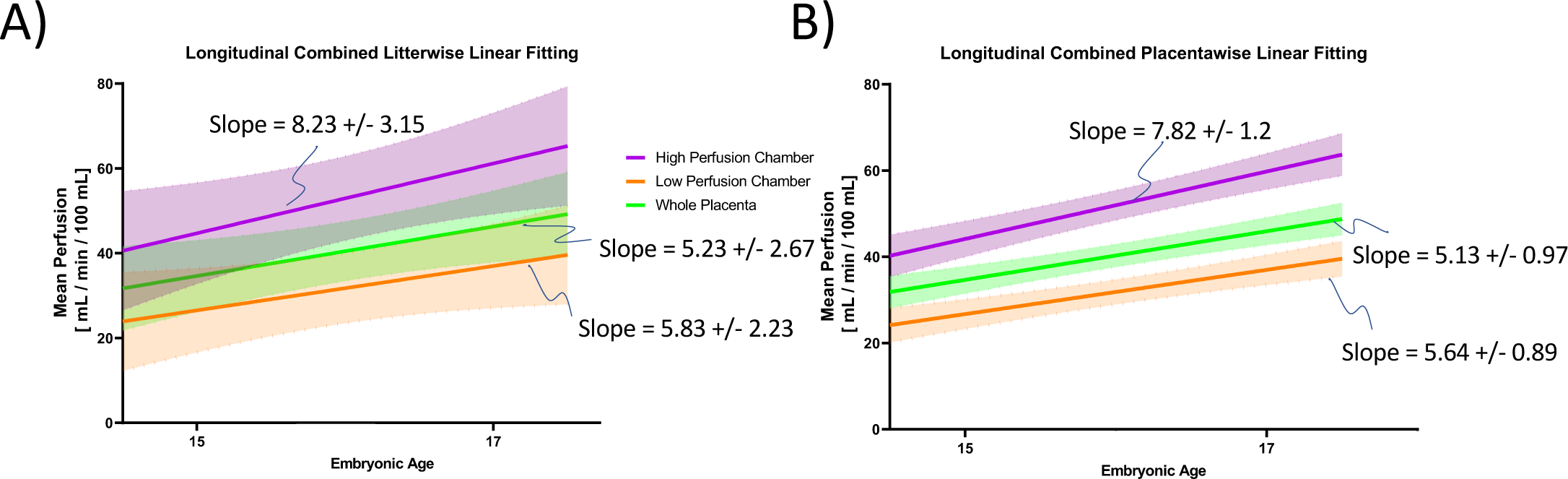
Longitudinal perfusion linear regression calculated per litter (A) and per placenta (B), split by the separate perfusion chambers. Seen in all plots is a linear increase in the mean perfusion across all chambers and the whole placenta from E14.5 to E17.5. Note that near complete separation of development trends at the placenta level are washed out when averaging across litters. Shaded areas represent 95% confidence intervals. Notice that in both the litter wise data and the placenta wise data, the slope of the trendline is much higher for the high perfusion chamber. The slope for the whole placenta nearly follows that of the low perfusion chamber.

### Probabilistic Quantification of Spatial Perfusion Patterns, Apparent Blood Volume, and Chamber Volume

In addition to placental perfusion and perfusion dynamics, we further developed novel biomarkers based on the probabilistic distribution of perfusion across the mouse placenta. We have successfully modeled the placental perfusion by a Gaussian-mixture-modeling to de-convolve two unique probability density functions (PDFs) from the original distribution of the log-transformed perfusion distribution. These PDFs allow further quantification of how perfusion is spatially distributed across the entire placenta. Figure 5A, B illustrates example PDF overlaid on the original histogram of the perfusion distribution for the same placentas on E14.5 compared to that of the ones on E17.5. The histograms of the placental perfusion can be fitted with two distinct Gaussian distributions (Fig.5AB) – the blue Gaussian distributions corresponding to the high perfusion compartments whereas the green Gaussian distributions corresponding to the low perfusion compartments. There is a noticeable increase in size and peak of the distribution positioned with a larger mean perfusion on E17.5 compared to that on E14.5, reflecting the increase in size of the high perfusion zone. This models the increase in size and vasculature of the labyrinth zone from E14.5 to E17.5. In addition, novel quantitative biomarkers can be extracted from PDFs for probabilistic quantification of placental development. Table 1 summarizes biomarkers isolated from the de-convolved Gaussian PDFs, describing the shape of the Gaussian distributions: the peak is the max height of each perfusion distribution, the half-width is the width of the distribution at half of the peak value, mu is the mean and sigma the standard deviation of the distribution, the area under the curve (AUC) is the trapezoidal numerical integration of each distribution function.

**Table 1).** Potential biomarkers derived from the probability density functions in **Figure 7**. The biggest differences from E14.5 to E17.5 can be seen in the peak low and high parameters as well as the AUC of the two distributions.

|  | **E14.5** | **E17.5** |
| --- | --- | --- |
| **Parameter** | **Mean ± SD** | **Mean ± SD** |
| Peak – Low Distribution | **1.63 ± 0.52** | **1.69 ± 0.44** |
| Peak – High Distribution | **1.20 ± 0.20** | **1.44 ± 0.32** |
| Half-Width – Low Distribution | **0.41 ± 0.12** | **0.35 ± 0.04** |
| Half-Width – High Distribution | **0.23 ± 0.04** | **0.26 ± 0.04** |
| Mu – Low Distribution | **1.54 ± 0.09** | **1.60 ± 0.06** |
| Mu – High Distribution | **1.55 ± 0.12** | **1.72 ± 0.12** |
| Sigma – Low Distribution | **0.04 ± 0.02** | **0.02 ± 0.01** |
| Sigma – High Distribution | **0.01 ± 0.01** | **0.01 ± 0.00** |
| AUC – Low Distribution | **0.68 ± 0.02** | **0.51 ± 0.12** |
| AUC – High Distribution | **0.31 ± 0.02** | **0.49 ± 0.12** |

The size of the PDF is proportional to the volume of the placenta and the amount of blood flowing through the placenta over the imaging period. Therefore, the apparent total blood volume (ABV) can be estimated by integration of the perfusion histograms (Fig.5C). The integration of the high perfusion histograms (Fig.5AB, blue) represents the ABV of the high perfusion chamber, whereas the integration of the low perfusion hisograms (Fig.5AB, green) represents the ABV of the low perfusion chamber. This method revealed statistically significant increase in ABV in the whole placenta, as well as the high and low perfusion chambers (Fig 5C). This result is driven both by the increase in overall placental perfusion from E14.5 to E17.5 (Fig. 3) as well as the increase in placental size from E14.5 to E17.5. Figure 5D, E summarizes placental volume findings from our model. There is a statistically significant increase in the average size of the whole placenta from E14.5 to E17.5 (Fig. 5D). Further, the volumetric ratios (Fig. 5E) of the high perfusion chamber to the low perfusion chamber show a statistically significant increase (p < 0.0001). The average size percentage of the low perfusion zone on E14.5 is 54.2% ± 13.1% and 45.8% ± 13.2% for the high perfusion chamber. We observed a similar shift as described by Coan et al (22) for the decidua and labyrinth zone, where the low perfusion chamber’s mean size percentage is 42.8% ± 14.0% at E17.5 whereas the high perfusion chamber’s mean size percentage was 57.2% ± 14.1%.

**Figure 5).**
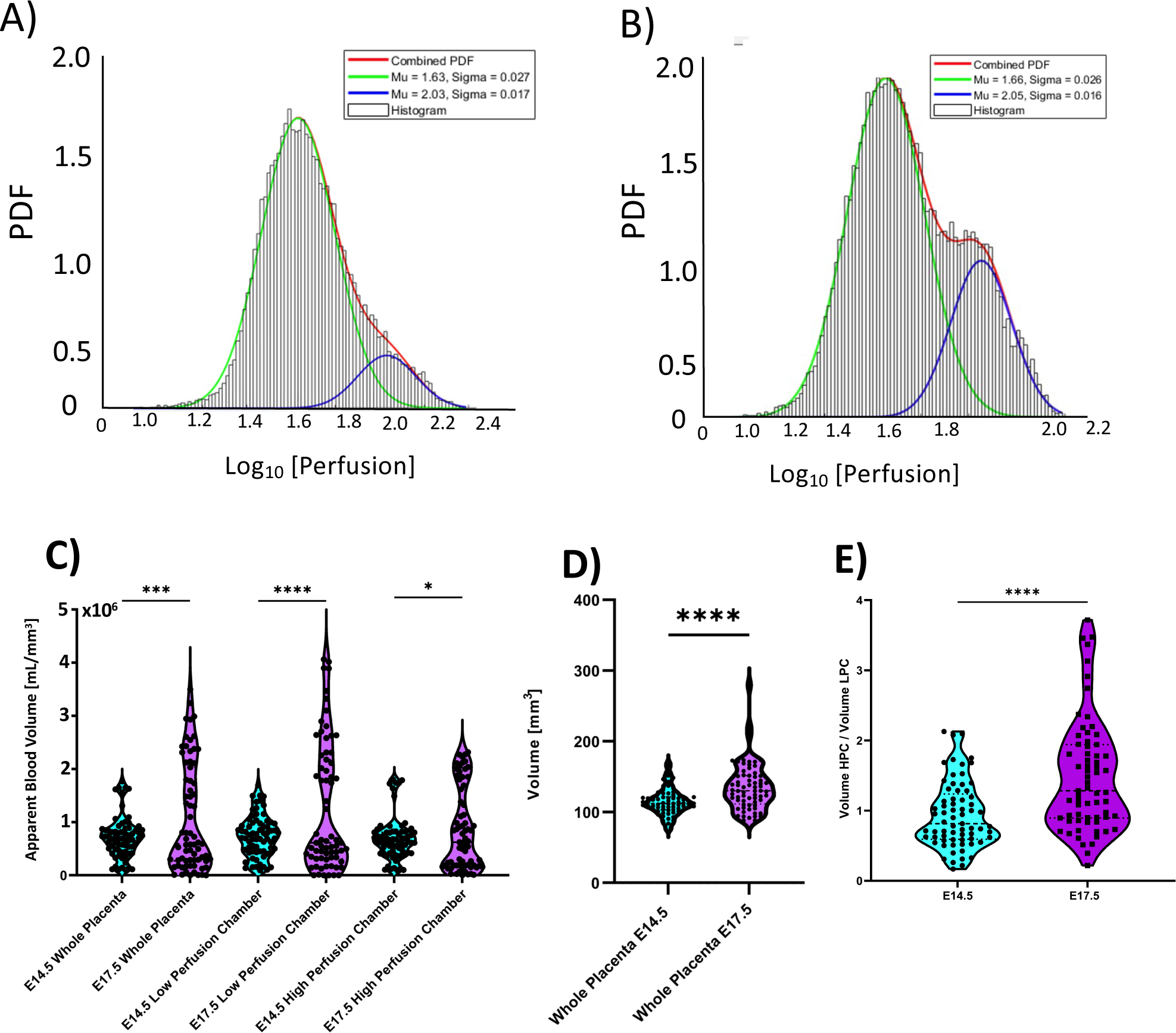
Probability density functions (PDFs) calculated from histogram estimations for the distribution of perfusion values within the whole mouse placenta on E14.5 (A) and E17.5 (B). The shape of the distribution suggests the presence of two separate PDFs, one for the two main perfusion chambers. The development from E14.5 to E17.5 demonstrates an increase in the size of the high perfusion distribution (blue PDF). The two separate distributions were successfully deconvolved and parametrized using Gaussian Mixture Modeling. Column plots of placental volume and blood volume development from E14.5 to E17.5. D) There is a statistically significant increase in the apparent blood volume in the whole placenta and both perfusion chambers from E14.5 and E17.5. B) Box and violin plot illustrating the slight, yet statistically significant increase in placental volume from E14.5 to E17.5. E) Lastly, there is a statistically significant increase in the ratio of the high perfusion chamber to the low perfusion chamber from E14.5 to E17.5. This nicely illustrates the selective increase in chamber size for the high perfusion region from E14.5 to E17.5. (* = p < 0.05, *** = p < 0.001, **** = p < 0.0001).

### Spatial Distribution of Apparent Gd Uptake Correlates with Perfusion MRI

We furthered our physiological-based chamber model with the longitudinal relaxation rate R_1_ which is the inverse of the longitudinal relaxation constant T_1_. The change in tissue R_1_ is proportional to the Gd accumulation. Thus, changes in placenta R_1_ due to Gd accumulation reflects the substrate delivery and accumulation in the placenta over the imaging period (∼900 seconds). Figure 6 (A-D) illustrates R_1_ maps on E14.5 (Fig.6A, C) and on E17.5 (Fig.6B, D) before (Fig. 6A, B) or after (Fig. 6C, D) physiological thresholding. We have successfully distinguished the high and low compartments in the quantitative R_1_ space. R_1_ maps revealed spatially distinguished compartments of similar sizes and locations as the high and low perfusion chambers delineated by the steepest slope (Fig.2) and perfusion (Fig.3) maps. Longitudinally, there is a statistically significant increase in the apparent Gd taken up by the whole placenta (Fig. 6E) (p = 0.247), the low (p < 0.0001) and the high perfusion chambers (p < 0.001) from E14.5 to E17.5 (Fig 6E). These findings match the trends seen in the changes in average apparent peak perfusion.

**Figure 6).**
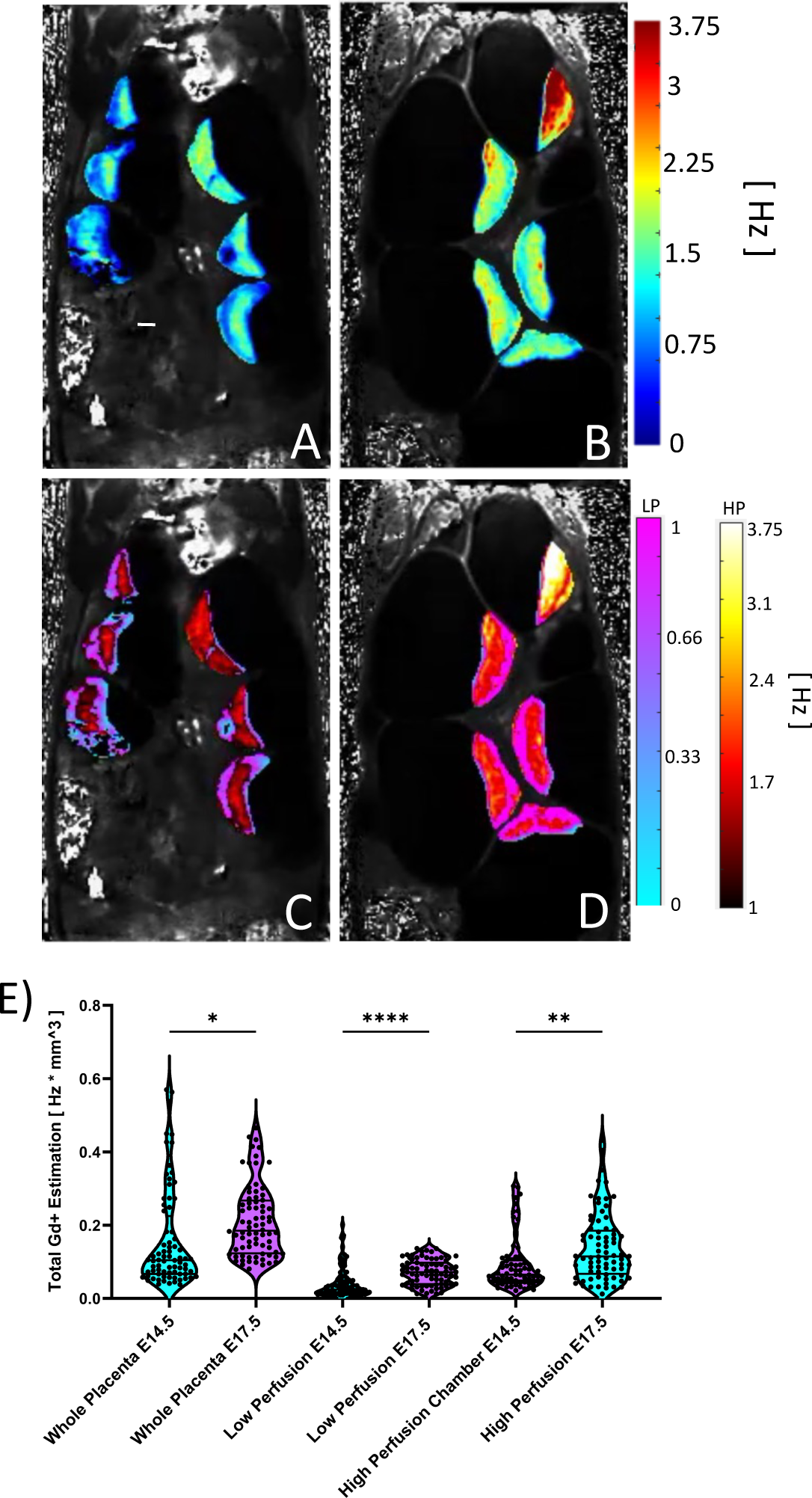
A and B Quantitative R_1_ maps prior to physiological thresholding for A) E14.5 and B) E17.5. C and D) the same R_1_ maps after physiological thresholding for C) E14.5 and D) E17.5. Physiological thresholding is again able to deconvolve different chambers in the placenta in the R_1_ space. The change in R_1_ is calculated pre-and post-administration of contrast agent and the apparent Gd uptake is estimated after normalizing by placental-chamber volume. E) Violin plots for of the average apparent Gd uptake for each placenta, separated by chamber as well as embryonic age. There is a statistically significant increase in the Gd uptake for the whole placentas as well as the low perfusion chamber from E14.5 to E17.5. This is not seen in the high perfusion chamber. (* = p < 0.05, **** = p < 0.0001). Scale bar is two centimeters.

### Characterization of Spontaneous Uterine Contractions

Time-series of the DCE images revealed extensive regular and phasic uterine motions for some placenta-fetal units. This phasic motion was still present even after removing maternal respiration motions with temporal median filtering and eliminating random large translations between frames with morphological closing. This motion was localized to the placenta-fetal units and appeared to be more pronounced on E17.5. Visual examination of each placenta revealed that the phasic motion was present in ∼11.1% of the placenta on E14.5, versus ∼82.1% on E17.5. Optical flow quantification allows us to visualize this motion in the form of velocity vectors overlaid on the DCE-MRI time series. Figure 7A, B shows continuous time frames of a single slice of DCE-MR images with the optical flow velocity vectors overlaid on E14.5 (Fig.7A) and E17.5 (Fig.7B). Larger magnitude of velocity vectors were seen in the E17.5 time series compared to those of the E14.5 ones. The averaged magnitude of the velocity vectors were significantly larger on E17.5 than on E14.5, regardless if the motions were measured from the high perfusion chamber, the lower perfusion chambers, or across the entire placentas. (Fig.7C). This indicates that the uterine motions were more profound with larger amplitudes on E17.5 than on E14.5.

**Figure 7).**
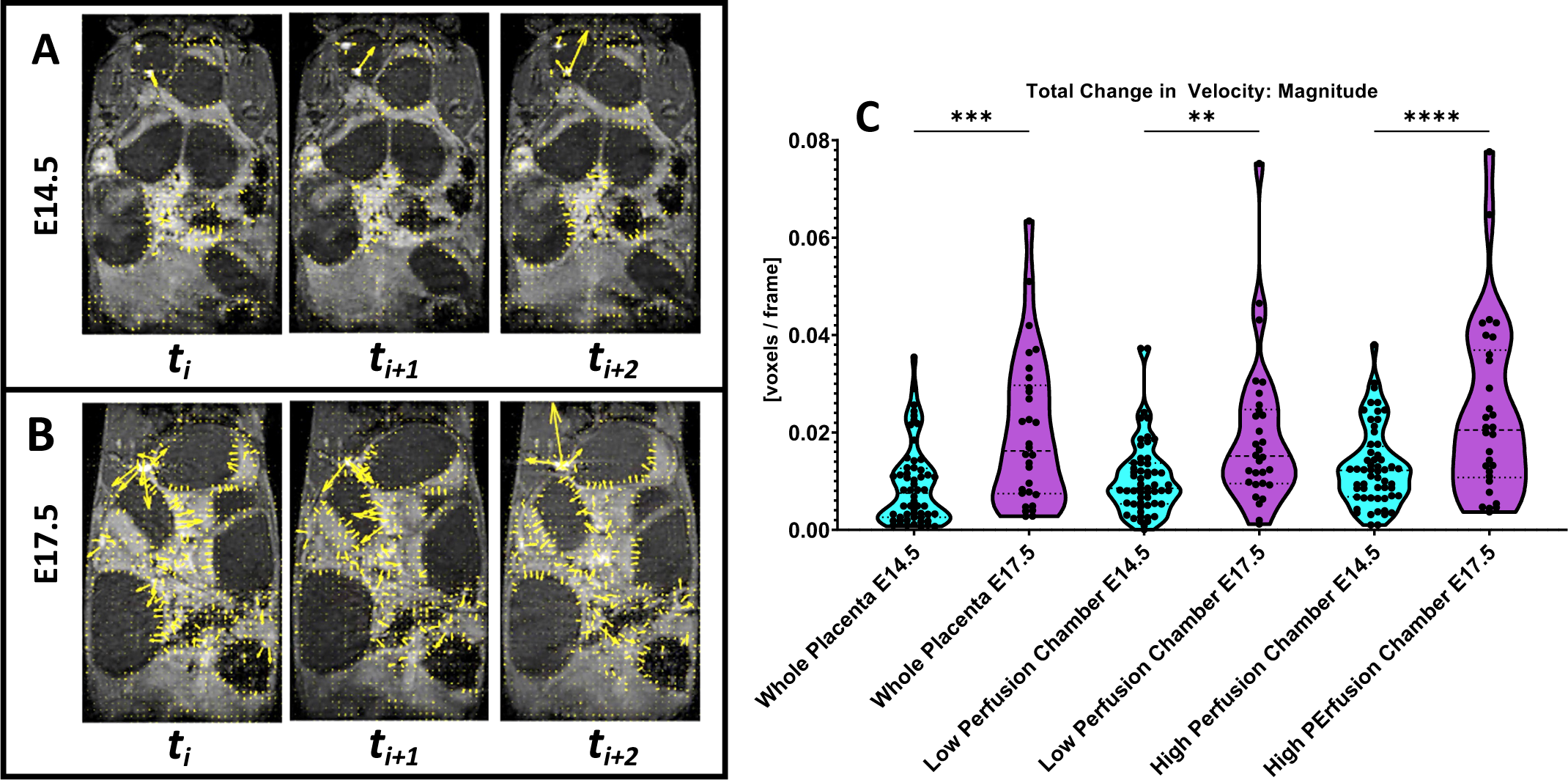
Results of optical flow apparent motion analysis. Flow vectors are overlaid on the original MR images for a single slice of the same animal at A) E14.5 and B) E17.5 over continuous frames. Average magnitude of velocity vectors for the whole placenta and its different perfusion chambers are compared via violin plot in C. (** = p < 0.01, *** = p < 0.001, **** = p < 0.0001)

Time-frequency and power analysis of the phasic motion traces for each placenta (Fig. 8) and its chambers revealed differential frequency components on E14.5 (Fig. 8 A, B) compared to those on E17.5 (Fig. 8 C, D). Periodograms (Fig.8 A, C) were used to identify the dominant frequencies in the time series, whereas the spectrograms (Fig.8 B, D) were used to express the power of the dominant frequencies of these phasic uterine motions over the imaging session. On E14.5 (Fig.8 A), the periodograms expressed the highest power only as the frequency approaching zero, illustrating the strength of the mean of the signal and is normal for most time series with some autocorrelation. The spectrogram of phasic motions on E14.5 (Fig. 8B) demonstrated that there were no instantaneous changes in frequency components over the time course. Further, when this data is stratified by the different perfusion chambers for the high perfusion chamber (Fig.8AB, left), lower perfusion chamber (Fig.8AB, middle) or the whole placenta (Fig.8 AB, right), no differences appeared in the periodogram estimation or the spectrograms among different compartments. On the other hand, looking at the phasic motion signals from E17.5 (Fig. 8C, D), specifically the periodograms (Fig.8C) and spectrograms (Fig.8D) for the high perfusion chamber (Fig. 8CD, left) exhibited an increase in frequency components from around 8-12 mHz, 15 mHz, and 25 mHz. However, the frequency characterization for the low perfusion chamber (Fig.8CD, middle) matched with that of all chambers on E14.5 (Fig.8 AB). The periodograms (Fig.8C, right) and spectrograms (Fig.8D, right) of the whole placenta on E17.5 appeared as an average of the frequency elements contributing from the high (Fig.8CD, left) and low perfusion (Fig.8 CD, middle) chambers, representing asymmetric motion of the mouse placenta on E17.5. All these frequency elements from the phasic motions are much slower than the maternal respiration (∼1.3 Hz) or heart rates (∼6.7 Hz).

**Figure 8).**
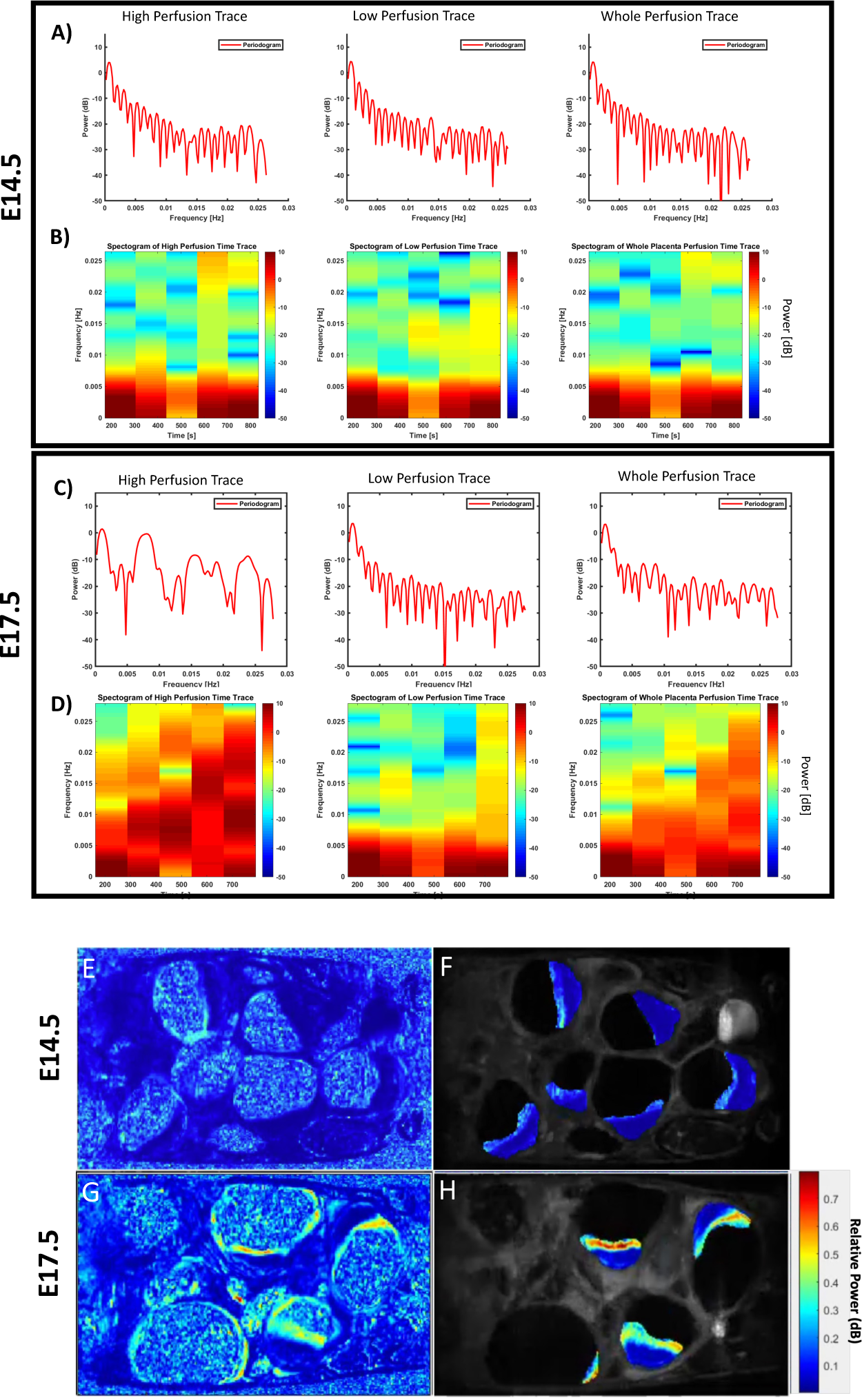
Power Spectral estimation and Time-Frequency analysis of the apparent motion time courses. A) PSD estimation and B) spectrogram at E14.5, C) PSD estimation and D) spectrogram at E17.5. Notice the lack of power present in the higher frequencies at E14.5 to E17.5 in the high perfusion chamber. This change in frequency power was not seen in the low perfusion chamber and seen in the whole perfusion chamber because of whole placental averaging. Relative Spectral Density sub band maps for the same litter at E-F) E14.5 and G-H) E17.5. E and G are the same images as F and H except with placentas segmented and overlaid on top of their original DCE-MR frames. E and G are provided to provide context of the relative power seen throughout the rest of the imaged tissue. Note that much of the spectral density is localized to the same region, with the same orientation, as the high perfusion chamber.

We further investigated the spatial localization of the most dominant sub-bands from the power spectrum analysis. The relative power spectrum of the 8-12 mHz sub band (Fig. 8E-H) was computed as a ratio to the total power in the entire spectrum range for each voxel in the DCE-time series and viewed as relative power maps for E14.5 (Fig. 8E) and E17.5 (Fig. 8G). These relative power maps demonstrated that the voxels showing the 8-12 mHz phasic motions were localized to the placenta-fetal units, but not the maternal abdominal organs. No relative frequency components within the 8-12 mHz sub bands were present in the kidney or across the abdomen but were specifically localized to the placenta and uterine walls. Moreover, the voxels with an increase in spectral density across the 8-12 mHz sub bands were localized to the high perfusion zones in the placenta. We confirmed this further by applying the placental segmentations to the relative motion power maps, and then overlaid them over the original MR anatomical images (Fig 8F, H). The voxels with the highest relative power of the phasic motions coincide with the voxels with the high perfusion chambers. The relative power maps of the phasic motions (Fig. 8 F, H) displayed the same spatial pattern as the high perfusion chambers seen in the dynamic perfusion characterization of the different placental chambers (Figs. 2,3).

## Discussion

### Physiological Thresholding demonstrates Spatial Variance of Placental Chambers across Multiple Perfusion MRI Domains

Although the identification of spatially varying perfusion zones within the mouse placenta has been identified in other studies (38–42, 44, 45), we expanded this knowledge with the addition of a biologically based (22) modeling for automatic localization of the different perfusion chambers. All other previous studies rely on manual analysis and region of interest (ROI) generation after data review. Instead, our group used a numerical threshold of the perfusion probability density function to localize the different perfusion chambers. We have presented a successful physiological model of placental perfusion development and motion dynamics for the mouse placenta by expanding on steepest-slope perfusion modeling and utilizing a physio-anatomical foundation for modeling the different placental chambers in quantitative perfusion MRI studies. Our model of the placental chambers and quantification of their physiological differences follows the expected development based on biomolecular studies of decidua and labyrinth formation (46). This is the first study of placental chamber formation that crosses from the quantitative perfusion space to other metrics of MR imaging. The same physio-anatomical thresholding method was utilized in the steepest-slope, apparent perfusion, and the quantitative R_1_ spaces. As the steepest-slope, average apparent perfusion, and R_1_ space elicit different meanings: instantaneous perfusion rate, average apparent peak perfusion, and local apparent gadolinium uptake, respectively. Specifically, changes in AGU demonstrate that the “wash-in” and “wash-out” mechanisms of the mouse placenta show differential kinetics across the various perfusion chambers and across embryonic age. Models of placental disease may elicit differences in one or more, or a combination thereof the different perfusion metrics within each perfusion chamber.

Further, contributions from each de-convolved perfusion PDF follows the proportions defined within our physiological thresholding method. This method combined with a model for the automated segmentation of the placenta within the DCE time-series would provide a full pipeline for automated chamber perfusion analysis.

An important characteristic of the methods and models developed in this paper is the non-invasive nature of image acquisition. The ability to non-invasively estimate important physiological parameters, such as mean apparent perfusion and apparent total blood volume, provides robust and quantitative biomarkers for longitudinal investigation of placental development of multi-fetal mouse pregnancy to complement the molecular and mechanistic investigation in preclinical mouse models. It also has a higher potential for clinical translation of the described methods. Many development associated diseases and disorders lack biomarkers that can effectively predict the variation of disease outcomes (6, 47–50). In addition, many congenital disorders and placental related conditions showed incomplete penetrance and variable outcomes. The underlying mechanisms for the variable phenotypical presentations cannot be understood without sensitive differential phenotyping tools. This major knowledge gap can be addressed by combining longitudinal functional MRI methods with invasive, molecular investigations. Such an investigation has the potential to drastically increase our understanding of developmental disorders, and our methods serve as a crucial support.

Future work will employ quantitative methods to correctly identify each placenta for successful longitudinal mouse imaging with multi-feal pregnancy. Mouse uterine horns are doubly perfused by the uterine branch of the ovarian artery and the distal part of the uterine artery (51, 52). A bi-directional arterial spin labeling (BD-ASL) approach (52, 53) can determine fetal/placental positions along each uterine horn. By saturating the proton spins in the incoming arterial blood in the uterine and ovary arteries, which dually perfuse each placenta-fetal unit, changes in placental signals correlated with its position along the uterine horn.

### Importance of the Placental Perfusion Development Program

The dramatic changes in placental function from E14.5 to E17.5 further illustrate how the placenta should be an important consideration when studying fetal physiology and neurodevelopment. Studies that do not take the time to consider the different contributions of each placental chamber therefore may not consider the full picture. The dramatic changes in the labyrinth zone as modeled by the high perfusion chamber have the potential to be vital to overall fetal development, in particular, neurodevelopment. In the future, we will use this model to quantify placental perfusion dynamics in various neurodevelopmental disorders, with the hypothesis that alterations in labyrinth zone perfusion dynamics are present in abnormal developing fetuses. We expect to see alterations in the default development program described in this study. If placental development is interrupted, the increase in perfusion, spatial patterning of perfusion, and/or end-term contractions may all demonstrate different dynamics with the potential of shining light on unanswered questions in fetal development diseases and disorders.

### Uterine Contractile Motion Effect on Analysis

The skew of contractile motion present at E17.5 compared E14.5 behooves the idea that this phenomenon may be a result of late-term contractions associated with pregnancy. Embryonic age 17.5 in the mouse is analogous to the late third trimester in human pregnancy (54, 55), where spontaneous Braxton-Hick’s contractions are often reported (30–32). Further, Gravina et al, (36) performed an extensive electrophysiological study into cervical, vaginal, and uterine tissues isolated from late-pregnant mice (E17 to E19) and found spontaneous, phasic contractions in all. Additionally, Nyquist sampling theory tells us that the highest frequency we can resolve without aliasing is half the temporal sampling frequency. The average respiration rate of anesthetized adult mice is ∼80 breaths per minute or ∼1.3 Hz, whereas the average heart rate of an anesthetized mouse is ∼400 beats per minute or ∼6.7 Hz. The temporal sampling frequency of our DCE-MRI protocol is ∼18 seconds, or 55.6 mHz. Thus, the highest frequency in the motion time courses we can resolve is half of our sampling rate at 27.8 mHz. This is drastically smaller than the frequency of anesthetized adult mice respiration and heart rate, 27.8 mHz vs 1.3 and 6.7 Hz, respectively.

One observation from our *in utero* contractile analysis was the asymmetric nature of apparent contractions. Increases in average velocity and changes in spectral density and time-frequency power heavily indicate a spatial patterning of contractile movement. Our observation of spatial patterning could be a result of spatially inhomogeneous contractions of the placenta or that the contractions are from a different tissue source and the spatial inhomogeneity seen in the placenta is a result of differential contractile propagation across differing uterine tissues. In future studies we will consider other uterine tissues and utilize signal processing tools to localize the origin of apparent uterine contraction to further characterize our observation.

The observed contractile phenomenon has variable effects across this entire analysis and our model. The steepest-slope modeling itself does not result in time-series perfusion maps, rather a temporal projection from a single frame where the steepest-slope of the imaging signal is calculated for every voxel of every slice in the time-series. However, if extreme motion is present, this could result in misleadingly high steepest-slope values because of placental contrast moving in and out of the imaging plane. Therefore, prior to determination of the steepest-slope map from the kinetic time-series data, the DCE time series were motion-corrected and spatially median filtered. As a result, the projection and estimation of the perfusion is not based on steady-state averages, but dynamics calculated for a single time, meaning that motion across the entire time-series does not play a role in steepest-slope and perfusion estimation. Therefore, after applying simple motion correction through spatial median filtering, the parameters of this model including steepest-slope quantification, mean perfusion, and probability density function estimation are unaffected by the uterine contractions.

Perfusion trace waveform and placental volume are two metrics that must be specially handled because of uterine contractions. To prevent incorrect morphology of the perfusion trace for the placenta and corresponding chamber, perfusion traces were developed using full time series segmentations, where each perfusion time point was estimated by averaging the intensity values across the placenta and then across the different placental chambers. This difference in analysis methods was chosen to minimize the introduction of low-frequency filtering artifacts to the perfusion traces, which could alter perfusion trace metrics such as apparent blood volume estimation. Further, the magnitude of artifacts that may be introduced to the perfusion curves are much smaller than the magnitude of the steepest slope. Thus, the perfusion curves were estimated slice-wise by averaging across a given segmentation at a given time for the entire time-series datasets. This observation and corresponding correction are crucial to consider for implementing this model for other studies as the motion on E17.5 may result in incorrect interpretations of physiological function.

### Conclusion

In summary, we have developed an in-vivo pipeline for the longitudinal analysis of placental perfusion and uterine-related contractions. We demonstrate the capabilities of the steepest-slope modeling method for peak perfusion estimation from subQ Gd contrast agent that appropriates dam health over multiple injections. We expand the perfusion chamber paradigm for the mouse placenta via novel use of a physiologically based threshold for delineation of the separate systems. We apply this methodology to two metrics not previously reported, the steepest slope of the perfusion time course, and the apparent Gd uptake. The latter of which is a function of the change in longitudinal relaxation pre-and post Gd administration. We have shown that the placenta’s anatomical chambers are defined not just by peak perfusion, but also the instantaneous change in perfusion and total Gd contribution from the vascular system.

Notably our application of computer vision and digital signal processing are the first to quantify contraction-related motion non-invasively, in-vivo. We use these tools to show that the mouse placenta exhibits asymmetric motion due to in-utero contractions, with most of the motion localized to the high perfusion chamber. We have expanded the researcher’s toolbox to investigate these contractions in combination with drugs known to modulate pregnancy hormones, contractions, and labor.

## Methods

### Animals

All mice used in the study were inbred C57BL/6J wild-type (WT) mice obtained from Jax Lab (Jax Stock No. 000664). Mice were housed in the AALAS certified animal vivarium in the Rangos Research Center. Mice were provided with food and water ad libitum and kept on a 12:12 hour dark/light cycle. Once weaned at 28 days, male or female littermates were housed with 2-4 mice per cage until 3 and a half months of age for experiments. Timed pregnancy was determined by the presence of vaginal plugs after crossing a male and a female in the same cage overnight. Ten pregnant females with 70 placentas were examined. Bodyweights of the pregnant females on E14.5 and E17.5 were 28.32 ± 1.58 g and 33.16 ± 1.69 g respectively. All animals received humane care in compliance with the Guide for the Care and Use of Laboratory Animals^1^. All experiments were conducted with the approval by the University of Pittsburgh Institutional Animal Care and Use Committee.

### Anesthesia for in vivo MRI

All mice received general inhalation anesthesia with Isoflurane for *in vivo* imaging. The mice were placed into a clear Plexiglas anesthesia induction box that allows unimpeded visual monitoring of the animals. Induction was achieved by administration of 3% Isoflurane mixed with oxygen for a few minutes. Depth of anesthesia was monitored by toe reflex (extension of limbs, spine positioning) and respiration rate. Once the plane of anesthesia was established, it was maintained with 1-2 % Isoflurane in oxygen via a designated nose cone and the mouse was transferred to the designated animal bed for imaging. Respiration was monitored using a pneumatic sensor placed between the animal bed and the mouse’s abdomen while rectal temperature was measured with a fiber optic sensor and maintained with feedback-controlled warm air source (SA Instruments, Stony Brook, NY, USA).

### Single bolus Gadolinium (Gd) contrast agent administration

A 24-gauge sterile subcutaneous catheter (McKesson Medical-Surgical Inc. Richmond, VA) was inserted into the dorsal subcutaneous space between the back skin and the muscle. A single bolus of Gd (0.1 mmol/kg bodyweight, using 1:10 dilution of Multihance, Gadobenate dimeglumine injection solution, 529 mg/ml, Bracco Diagnostics, Inc. Monroe Twp, NJ) was administered between time frames 4 and 5 via the subcutaneous catheter during the dynamic contrast enhancement (DCE) MRI acquisition.

### *In vivo* MRI Acquisition

*In vivo* MRI was carried out on a Bruker BioSpec 70/30 USR spectrometer (Bruker BioSpin MRI, Billerica, MA, USA) operating at 7-Tesla field strength, equipped with an actively shielded gradient system, using a quadrature radio-frequency volume coil with an inner-diameter of 35 mm for transmission and reception.

For dynamic contrast enhancement (DCE), multi-planar T_1_-weighted dynamic time series was acquired using free-breathing-no-gating IntraGate fast gradient echo sequence with retrospective motion deconvolution using the following parameters: field of view (FOV) = 4.0 cm X 2.5 cm, matrix = 256 X 160, slice thickness = 0.85 mm, in-plane resolution = 0.156 mm, echo time (TE) = 1.872 msec, repetition time (TR) = 101.493 msec, flip angle (FA) = 75^0^, number of repetition (NR) = 1, acquisition time = 10 sec 120 msec.

R_1_ mapping was achieved by varied multiple flip angles. Five multi-planar IntraGate image stacks were acquired using the following parameters: field of view (FOV) = 4.0 cm X 2.5 cm, matrix = 256 X 160, slice thickness = 0.85 mm, in-plane resolution = 0.156 mm, echo time (TE) = 1.872 msec, repetition time (TR) = 114.88 msec, 5 flip angle (FA) = 75^0^, 55^0^, 30^0^, 20^0^, and 10^0^, number of repetition (NR) = 10, acquisition time = 1 min 33 sec 904 msec.

### Image Pre-Processing and Segmentation

Prior to quantitative analysis, temporal morphological closing was performed to remove any potential motion artifacts. Spatial median filtering was applied to each image in the time-series for further removal of motion artifacts and smoothing of high-frequency noise. At this point steepest-slope modeling was performed. Finally, a mean-removed maximum intensity projection (MIP) was created from the DCE time-series for each slice. The mean kinetic image was calculated by averaging MR images acquired during the initial uptake phase of the administered gadolinium. The result was a 256 by 160 by 20 slice image-matrix. This matrix was output to ITK-snap where manual segmentation of each placenta in the litter was performed. The segmentations were masked on the perfusion maps of each animal, and all quantification was then performed.

Kinetic time series of each slice were also created, resulting in a 256 by 160 by 50 time series image-matrix. These matrices were exported to ITK-snap where manual segmentation of the placenta was performed. These secondary masks were used to produce all time-series traces. The final quantitative R_1_ maps pre-and post-contrast agent were also output to ITK-snap where manual segmentation was again performed. These segmentations were used for R_1_ and Gd metrics as well as to calculate placental volume.

### Quantitative R_1_ Mapping and Apparent Gd Quantification

Gadolinium-based contrast agents alter the longitudinal relaxation time associated with proton spin that is responsible for the MRI signal (56, 57), this longitudinal relaxation time is also known as T_1_. The paramagnetic properties of Gd decrease the T_1_ relaxation time for a given tissue. The inverse of T_1_ is known as R_1_ and is measured in Hz, with an increase in tissue R_1_ being directly proportional to the apparent amount of Gd taken up by the tissue of interest.

Similar metrics looking at changes in tissue relaxation times because of Gd uptake can be seen in studies that quantify extracellular volume by scaling changes in R_1_ for a specific tissue by the subject’s hematocrit level (57–59). Changes in placenta R_1_ due to Gd accumulation reflects the substrate delivery and accumulation in the placenta. Therefore, determining the change in R_1_ pre-and post-contrast can be used to further understand development of material-transfer sysems in placental tissue. To determine the total apparent Gd placental uptake, T_1_ relaxation quantification was carried out for each litter pre-and post-administration of the contrast agent using the varied-flip angle method (60). The varied flip-angle method utilizes the relationship between the steady-state MR signal *S_MR_* and T_1_/TR as described by the Ernst Equation:

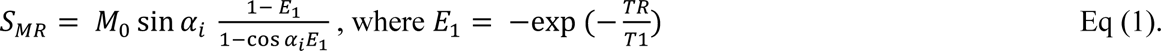

This equation is linearized with slope *E_1_* and T_1_ can be extracted from the slope via the following:

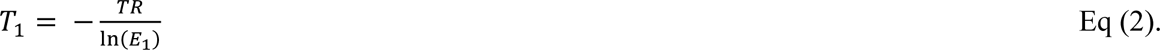

The slope, E_1_ is numerically determined from the Intragate FLASH image data via a least-squares based nonlinear fitting of equation 1. Intragate FLASH imaging was utilized with flip angles of 75, 55, 30, 20, and 10 degrees. Imaging parameters included TE/TR = 1.872/114.8 ms with 10 repetitions. The field of view was set to 40 x 25 mm with a matrix of 256 x 160 for a final resolution of 0.156 mm/pixel by 0.156mm/pixel. Varied Flip Angle imaging was performed directly prior to subcutaneous administration of the contrast agent and directly after perfusion kinetic imaging was completed.

Finally, the full-tissue apparent Gd uptake (AGU) was estimated using the following equation that relates the change in R_1_ pre-and post-contrast to the volume of the placenta and its corresponding chambers.

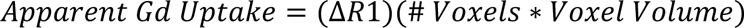

### Perfusion Estimation via the Steepest Slope Model

The steepest slope of a given voxel intensity gradient represents a faster increase in local concentrations of administered contrast agent. The steepest slope modelling schema is a one-parameter model to estimate instantaneous tissue perfusion using DCE-MRI. Brix et al. (38) provide a comprehensive review of the steepest slope model as a tool for tissue perfusion estimation and clinical efficacy. The tissue concentration of Gd during initial wash-in is represented by the following equation:

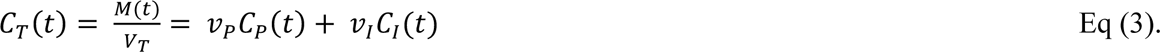

Where C_T_ is the Gd tissue concentration, M(t) is the mass of the Gd in the placental volume. V_P_ and V_I_ represent the ratios of intravascular and interstitial spaces of the placenta, respectively. Finally, C_P_ and C_I_ are the Gd tissue concentrations in the respective intravascular and interstitial spaces.

Further, assuming Gd travels in the blood with plasma flow F, then the mass gradient over time of Gd within the placenta can be described by Fick’s Principle (61). Fick’s principle states that the calculation for blood flow to a given tissue can be calculated if the Gd concentration at the arterial tissue input, the venous tissue output, and the amount taken up by the organ is known. Incorporating plasma flow F, the mass-gradient of the Gd over time becomes:

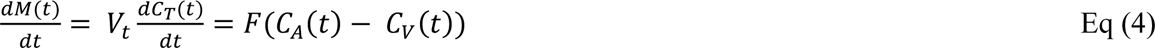

Where C_A_ is the tissue arterial input function (AIF), and C_V_ is the venous output. Finally, by assuming that no venous wash-out occurs during uptake of Gd, the concentration-gradient of the contrast agent over time becomes:

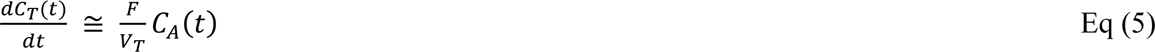

It can now be seen that Equation (5) provides a relationship between the perfusion in the tissue over time and the concentration of the administered Gd. Final perfusion value for each voxel is then calculated as the max perfusion of the voxel over time. Therefore Eq. (5) reaches a maximum when the left-hand side reaches its maxima.

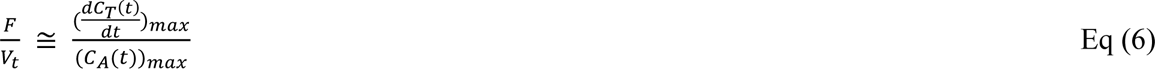

The arterial input to the placental chambers is taken from the kidney renal hilum of each litter mother. A small region is hand-annotated over the renal hilum and the corresponding AIF is produced from each DCE time-series MRI image. The change in contrast agent over time is estimated from the raw DCE time-series images for every voxel. The voxel-time point with the greatest estimated change is kept and divided by the max of the AIF, as described by Eq. 6. This process reduces the dimensionality of the time-series DCE to a single map of tissue perfusion. This process is repeated for every 2D-DCE slice taken to produce a final 3D perfusion map.

### Physiological Thresholding of the Mouse Placenta into Perfusion Chambers

The identification of separate perfusion chambers in the mouse placenta via DCE-based MRI has been shown in several other studies that correlate the perfusion chambers with the decidua and labyrinth (39–41). These zones are anatomically and physiologically defined and separated by a junction-protein dense membrane to regulate endocrine crosstalk between the two regions (54). A main function of the labyrinth zone is to mediate nutrient and waste exchange between the maternal blood and the amniotic fluid of the embryo (54). The labyrinth zone contains maternal blood sinusoids that interface with fetal blood vessels in the form of a dense network of vessels that utilize counter-current blood flow for rapid nutrient exchange. The extensive surface area of this network makes it a dense blood highway with increased perfusion compared to the decidua and junctional zone (54).

Coan et al (22), have performed an exhaustive stereology-based analysis of the volume of various placental chambers. They performed the analysis for E14.5, E16.5, and E18.5 litters. Estimation of E17.5 was performed by averaging the results for E16.5 and E18.5. They conclude that at E14.5, the labyrinth zone takes up roughly 45% of placental volume and 55% is taken by the decidua and junctional zone. These proportions switch on E17.5, however, and roughly 55% of the volume is taken up by the labyrinth zone. Therefore, automatic localization of each the different perfusion chambers in each segmented placenta is performed by calculating the 55^th^ and 45^th^ percentile of placental perfusion values on E14.5 and E17.5, respectively. Any voxels below this threshold are deemed in the ‘Low Perfusion Chamber’ and voxels above deemed in the ‘High Perfusion Chamber.’

### Perfusion Dynamics

The steepest slope-model’s single parameter is the peak of the gradient of concentration of the administered contrast agent through the placental chambers. After thresholding, this parameter was collected from each voxel from E14.5 to E17.5. The average perfusion value of the different placental chambers and whole placenta was also calculated for the different groups. Apparent total blood volume (ABV) through the placenta was estimated via the following equation:

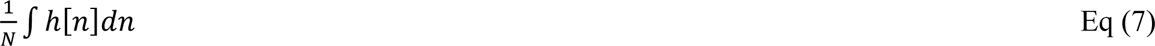

Where 1/N is the size of the placenta calculated as the number of voxels in the segmentation mask. H[n] is the probability density function of the placental perfusion. Apparent total BV was calculated for the high and low perfusion chambers after physiological thresholding using the same equation. This method was implemented numerically using trapezoidal estimation of the area under the perfusion trace.

### Numerical Modeling using Gaussian Mixture Modeling

The Probability Density Function (PDF) was first estimated using histogram functions in MATLAB. The PDF was estimated from the perfusion values for all voxels in the segmented placentas for each litter. The histograms were Log-base-10 transformed to achieve a more normal probability distribution. Then, a two-component Gaussian mixture model was used to estimate and parameterize the true PDFs from the histogram estimates. The Expectation-Maximization algorithm (62) (EM) was implemented via built-in MATLAB functions to estimate the probability parameters 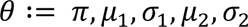_;_ of the perfusion distribution. The EM algorithm is a method for determining the maximum likelihood estimate (MLE) of the unknown probabilistic parameters. The EM algorithm maximizes the marginal likelihood of observed data:

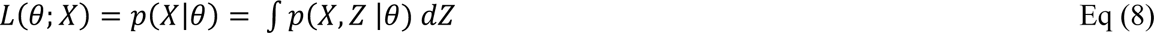

Where X is the observed perfusion values of each voxel in each placenta, Theta is a vector representing the PDF parameters mu and sigma, and Z is unobserved or missing values. As Z is generally indescribable, the EM algorithm solves this problem by first randomly initializing *θ*, and assigning each point to the two PDFs defined by its parameters. Then performing an *expectation* step by computing the responsibilities of each point to its corresponding PDF. *θ* is then updated via the *maximum-likelihood formulas.* The overall algorithm is described in detail below:

1. Random initialization of *θ*,
2. Expectation is performed via computation of the contribution of each data point to the two initial clusters:

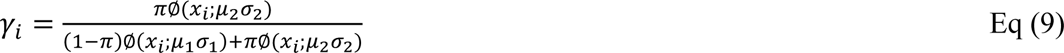
3. Maximization is performed and the parameters defined with *θ* are updated via weighting of the new parameters by *γ_i_*

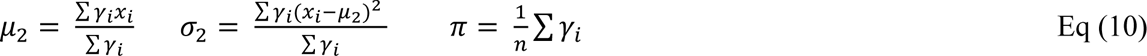
4. Steps two and three are repeated until convergence is reached.

After computing the parameter mu and sigma for each Gaussian component, features of the distributions were computed as potential biomarkers for placental development. For each distribution the following metrics were collected: area under the curve, the half-height, the full width at half maximum (FWHM) and peak value.

### Quantification of Placental Motion

Exploratory analysis of DCE time-series images revealed extensive placenta motion, specifically at E17.5, after normal motion correction was applied. This motion was quantified using a mathematical model of vision known as optical flow (63, 64). We utilize the Horn-Shuck (HS) method of solving the optic flow aperture problem to quantify the apparent motion across our DCE time series. The HS optic flow method was implemented to estimate placental motion in each slice of the DCE time-series images using built-in MATLAB functions. The regularization parameter that controls the expected smoothness of the input was set to 10^∧^2. The HS optical flow method results in a dense vector field that contains velocity vector information for every voxel at every time point. The time-series segmentations used to extract perfusion traces are also used to extract average velocity traces for each whole placenta, the low perfusion, and the high perfusion chambers. Average velocity magnitude for each traces was computed and compared across the whole placenta and corresponding perfusion chambers.

### Power Spectral Density Estimation and Time-Frequency Analysis of Placental Motion

The spectrogram and periodogram were both used to quantify frequency and spectral characteristics of uterine contraction velocity. The periodogram is an estimation of the power spectral density (PSD) of a given signal. In other words, the periodogram is an estimate of the frequency power per unit of frequency. Formally, it is mathematically in a continuous sense defined as the Fourier transform of the autocorrelation of the signal, as described by the following equation.

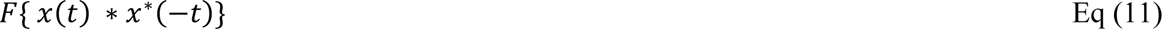

Where F is the Fourier transform operator and ∗ is the correlation operator. However, this definition cannot be used for discreate data and must be estimated using time-averaging. Various methods have been developed for this estimation, but in this study Welch’s method (65) is implemented using built-in MATLAB functions.

The spectrogram is a tool for performing time-frequency analysis that is, seeing how frequency power of a signal changes over the time course of the signal. The result is a two-dimensional matrix where each value in the matrix represents the power at a given frequency at a given time. In words, this is calculated by windowing a signal into segments with or without overlap and calculating the frequency power of that segment. The length of the window and amount of overlap controls the resolution in the time domain. In practice, this is implemented using a signal processing tool known as the Discrete short-time Fourier transform (Discrete STFT). The Discrete STFT can be calculated using the following equation:

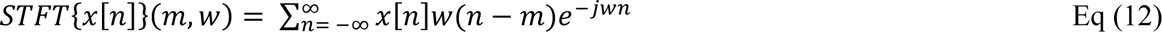

Further, the magnitude-squared of the equation above is formally known as the spectrogram:

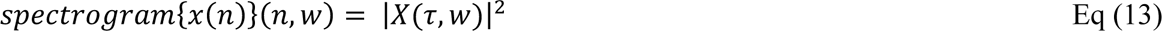

In this study, the above is implemented using built-in MATLAB functions. A 15-sample Bartlett window with 50% overlap between segments was used for the window, and the size of the fast-Fourier transform (FFT) was ten times the length of the DCE time-series.

Both above analytical efforts revealed the most frequency power and spectral density at and around the 10 mHz frequency band. Therefore, as our last effort into exploring this phenomenon, the periodogram was calculated for every voxel of the time-series images. Then, the relative power in the sub-band from 8 to 12 mHz was calculated. Then, the relative power spectrum of the 8-12 mHz sub band was computed as a ratio to the total power in the entire spectrum range for each voxel in the DCE-time series and view and as spectral density projection map of the time series.

## Statistics

Unpaired t-tests with unequal variances were performed for comparisons between two groups. One-way ANOVA was utilized for all other statistical analysis with Tukey-Kramer tests used for all ad-hoc testing. Statistical analysis performed using Prism 9 graphing and statistical software package.

## Study Approval

All animal protocols were approved by the Institutional Animal Care and Use Committee of the University of Pittsburgh.

## Author Contributions

DREC implemented and performed all image analysis, in charge of paper drafting and review, responsible for conception of physiological thresholding and conception of probabilistic quantification as well as time-frequency analysis. KES acquired MRI data. MCS and NWC were responsible for timing mouse pregnancy, colony management, and assisted with data analysis. DW performed ROI segmentation and analysis. MGO and RWP assisted with concept formation and study design. YLW is responsible for overall study design, conceptual formation, establishing protocols, training and supervising research personnel, data acquisition, and overseeing all aspects of the study.

## Acknowledgements

The authors wish to thank Ms. Caitlin Swanberg for participating in the preliminary testing phase. This work is partly supported by funding to YLW from NIH (EB023507, NS121706-01), AHA (18CDA34140024), and DoD (W81XWH1810070, W81XWH-22-1-0221).

